# The co-regulatory interaction of transpiration and water potential, and implications for stomatal conductance models

**DOI:** 10.1101/237826

**Authors:** Robert Beyer, Hans Pretzsch, Paul-Henry Cournède

## Abstract

Leaf water potential decreases with increasing transpiration rate according to an analogue of Ohm’s law, while transpiration rate decreases with decreasing leaf water potential in the framework of stomatal control. This interaction is not accommodated in present-day models of stomatal conductance. We formally derive the equilibrium between these two counteracting processes for steady-state water conditions. We show that the mechanism considered causes an attenuation of the immediate effect of atmospheric variables on transpiration, which can improve existing models of stomatal conductance that presume non-interdependent variables. Parameters from European beech (*Fagus sylvatica* L.) are used to illustrate the results.

## Introduction

Models of stomatal conductance *g_s_* at the leaf level commonly postulate a onesided dependence of *g_s_* on different environmental factors (Damour et al. 2010). The multiplicative approach of Jarvis (1976), expressing *g_s_* as a product of empirical functions of light intensity, leaf temperature, vapour pressure deficit, ambient carbon dioxide concentration and leaf water potential has had a lasting effect. In order to assess water stress more explicitly, follow-up work replaced the response to leaf water potential by a response to soil water deficit (Stewart 1988), instant pre-dawn leaf water potential (Misson et al. 2004) as well as predawn leaf water potential aggregated over a period of time (MacFarlane et al. 2004, White et al. 1999), the latter two variables being assumed a proxy for soil water potential.

The tendency of existing models to express stomatal conductance, *g_s_*, as a one-sided function of leaf water potential, ψ(ℓ), neglects the co-regulatory interaction between these two variables: Indeed, the response of *g_s_* to ψ(ℓ) is well documented for many species and conditions, and Damour et al. (2010) emphasise the relationship’s “mechanistic basis because it is well demonstrated that stomatal movements result from variations in leaf (or guard cell) water status, which result themselves from variations of evaporation in the substomatal cavity, and thus of the transpiration flux” (cf. also Buckley and Mott 2002). However, there is also a direct mechanistic impact of *g_s_* on ψ(ℓ). By analogy to Ohm’s law (see below), water potential along a soil-to-leaf water column is a function of leaf transpiration rate E, which is closely related to *g_s_* via

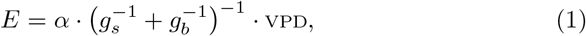

where *g_b_,* VPD and α denote boundary layer conductance to water vapour, vapor pressure deficit and a physical parameter, respectively (Damour et al. 2010). Hence, E decreases with decreasing ψ(ℓ), while ψ(ℓ) increases with decreasing E. In this article, we formally determine the equilibrium between these counteracting mechanisms. We show how the result can be used with existing models for stomatal conductance, and discusses implications for height growth in trees. We illustrate our findings using parameters of European beech (*Fagus sylvatica* L.).

## Model description

Denote by E ≥ 0 [kgs^−1^] the average transpiration rate from a leaf at a certain point in time. Let ℓ [m] denote the length of the soil-to-leaf pathway, and *K*(*s*) [kgs^−1^ m^−1^ Pa^−1^] the hydraulic conductivity of the root or shoot segment supplying the leaf and located *s* ∈ [0, ℓ] meters away from the soil-root interface. Under steady-state conditions, the water potential ψ(*s*) ≤ 0 [Pa] along the soil-to-leaf water column obeys

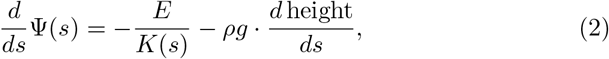

by analogy to Ohm’s law (Tyree and Zimmermann 2002). The last term is due to gravitational force, with *ρ* = 1000 kgm^−3^ and *g* = 9.81ms^−2^ denoting density of water and acceleration due to gravity, respectively. ψ(0) and ψ(ℓ) represent average soil and leaf water potential, respectively, at the given time. For leaf height *h* and given soil water potential ψ(0) = ψ_0_, (2) solves to

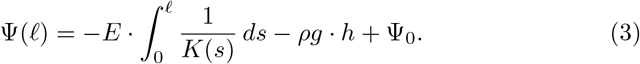

Tyree and Zimmermann (2002) established a sigmoid relationship of transpiration rate *E* against leaf water potential ψ(ℓ). *E* reaches its maximum for ψ(ℓ) = 0, and tends to 0 as ψ(ℓ) decreases. This corresponds to the closure of stomata in case of low leaf water potential in order to prevent the rupture of the root-to-leaf water column and the formation of embolisms (Tyree and Zimmermann 2002). Depending on whether this mechanism applies sooner or later, species are characterised as isohydric and anisohydric, respectively (McDowell et al. 2008). Let the response of *E* to ψ(ℓ) be described by an empirically obtained function

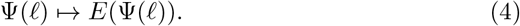

Inserting (4) into (3) results in an equation for ψ(ℓ),

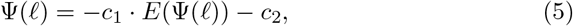

where for short 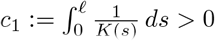 and c_2_:= *ρg · h* — ψ_0_ > 0.

According to (3), ψ(ℓ) decreases with increasing *E*, while *E* decreases with decreasing ψ(ℓ) according to (4). The solution of (5) represents the equilibrium between these counteracting mechanisms. It exists and is unique: The right-hand side has a unique fixpoint ψ(ℓ)^*^ < 0, since it is a monotonically decreasing function of ψ(ℓ), and negative at ψ(ℓ) = 0. The corresponding equilibrium transpiration rate follows from (4) as *E*^*^ = *E*(ψ(ℓ)^*^). Here, *E*^*^ varies only with soil-to-leaf hydraulic resistance 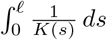, height *h* and soil water potential ψ_0_, which allows us to express it in terms of a species-specific function,

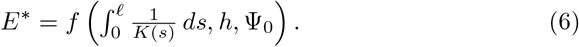

Equilibrium stomatal conductance 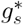 follows from (1). For suitable representations of the sigmoid function (4), *f* can be derived explicitly (Appendix A1).

A priori, the soil-to-leaf hydraulic resistance 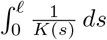 ds depends on individual tree anatomy. In Appendix A2, we argue that it may be approximated by a species-specific constant, in which case *E*^*^ could be expressed as a function of *h* and ψ_0_ alone.

### Generalisation to other environmental variables

Similar to Jarvis (1976) and follow-up work, the dependence of *E*^*^ (and analogously stomatal conductance) on soil water potential ψ_0_ can be readily extended to additional exogenous environmental variables such as radiation intensity, temperature, vapour pressure deficit and carbon dioxide concentration in a multiplicative way. (4) then generalises to

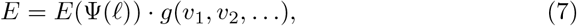

where 0 ≤ *g* ≤ 1 describes the relative response, i.e. rescaled such that it equals 1 at optimal *V_i_* values, of transpiration rate to the environmental variables *V_i_* at fixed ψ(ℓ) =0. The function g may, on its part, be the product of functions of one *V_i_* each, as done in many models. This extension affects the calculation in above section only in that the constant *c*_2_ in (5) changes. This is a consequence of the fact that the *V_i_*, unlike leaf water potential, are exogenous variables. We thus obtain a species-specific function, generalising (6):

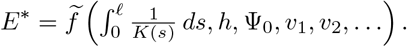

Unlike Jarvis (1976) and followers, this is not a multiplicative approach in the narrow sense, since varying the *V_i_* also affects ψ(ℓ) by inducing a different *E*^*^-ψ(ℓ)^*^ balance. If the immediate effect of varying *V_i_* is an increase (decrease) of transpiration rate, this increase (decrease) would be lessened since it leads to a small decrease (increase) in leaf water potential, which, on its turn, leads to a small decrease (increase) of transpiration rate. For instance, if *g* were linearly increasing (decreasing) in *v*_1_, then 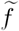 would be a concave (convex) function of *v*_1_. The response of *E*^*^ to varying *v*_*i*_ is thus more complex than the term *g* alone, specifically in that its immediate effect is attenuated as a consequence of the interaction of transpiration and leaf water potential.

### Simulation example

We illustrate the above results using parameters for European beech *(Fagus sylvatica* L.). We use the representation in Appendix A1 to describe the response of *E* to ψ(ℓ), for which parameters in (A1) were estimated as *p*_1_ = 1.7 ² 10^−4^, *p*_2_ = 1.8, *p*_3_ = 3, using data from Lemoine et al. (2002), and, for convenience, a linear approximation of (1), as suggested by Monteith (1995), *g_s_* ≈ 60.24. *E*, based on data from Cochard et al. (2000). 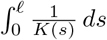 ds was set to 2.3 o 10^4^ (Aranda et al. 2005). To exemplify the property described in the above paragraph, we considered an additional, hypothetical environmental variable, *V_1_* ∈[0,1], for which the immediate response of *E* is assumed as *g*(*v_i_*) = *v_i_*, in the notation of (7). For *v_1_* ≡ 1, this reduces to the case described in (6).

Fig. 1 shows equilibrium leaf water potential and transpiration rate as a function of *v*_1_ as well as height and soil water potential, which were combined into one term. Several nonlinearities can be observed: As described above, even though *g* is linear, the response of *E*^*^ to varying *v_1_* is concave, i.e. the ultimate effect of varying v_1_ is attenuated. This would not be the case in a strictly multiplicative model. Similarly, ψ(ℓ)^*^ is a nonlinear function of *v*_1_. *E*^*^ shows a sigmoid response to –*ρg · h* + ψ_0_, which results from the qualitatively similar response of *E* to ψ(ℓ). Lastly, the response of ψ(ℓ)^*^ to changing ψ_0_ is nonlinear, which would not be the case if *E* were independent from ψ(ℓ) in (2).

**Figure 1:**
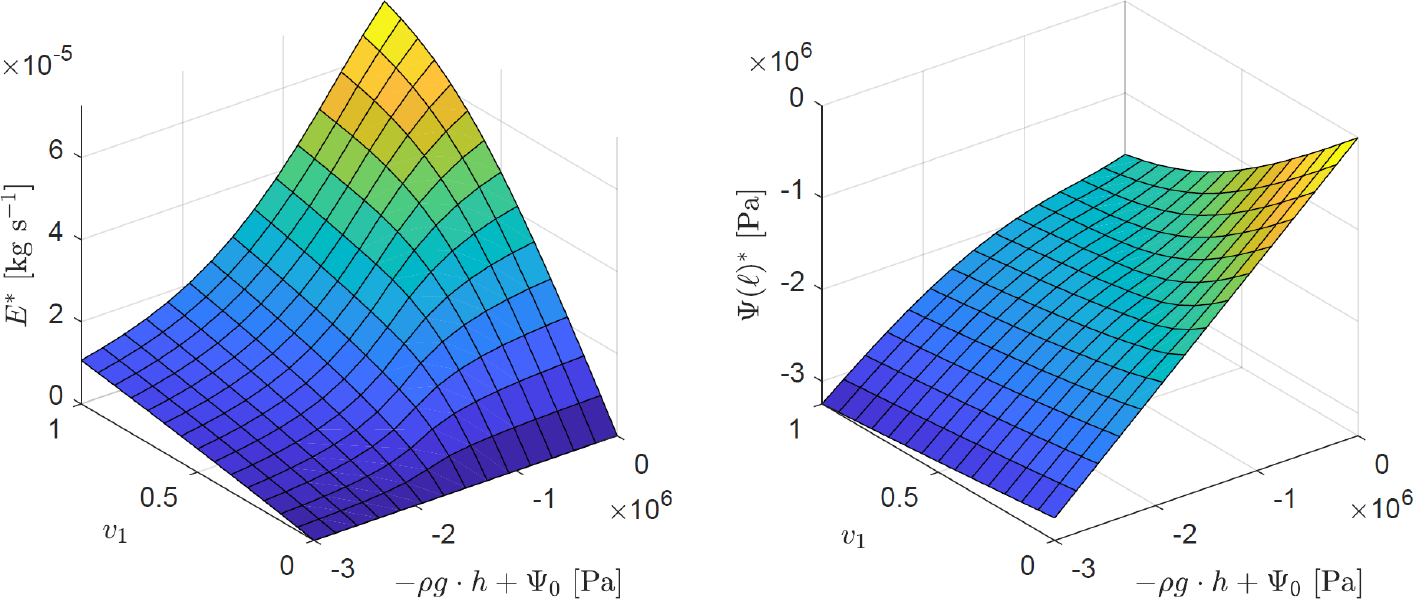
Equilibrium leaf water potential (left) and transpiration rate (right) in European beech as a function of two environmental variables (see text for details). Each additional 1 m in leaf height *h* corresponds to a reduction of —*ρg · h* + ψ_0_ by *ρg* = 9810 Pa.

## Discussion

The strong impacts that current climatic trends and weather events have on tree and forest growth require a sound understanding of the stress tolerance of trees (Hartmann 2011). Models can help to better map stress reactions and thus offer decision support for the selection of resistant and resilient species in order to avert future damage. The interaction of water potential and transpiration described here, as well as the demonstrated effect of this regulatory circuit on the impact of other environmental variables, contributes to this strategy.

We have shown that the relationship between leaf water potential and transpiration is not unilateral as tacitly assumed in current models, but interactive. We developed a model to accommodate this mechanism, and demonstrated how it can be used with existing models for stomatal conductance, in which it proved to attenuate the immediate effect of varying exogenous environmental variables. This addresses a major criticism of multiplicative approaches, namely the supposedly independent action of the different factors (Damour et al. 2010).

In (3), the roles of soil water potential, ψ_0_, and height, in terms of *ρg* ⋅ *h*, are interchangeable (cf. Fig. 1). Provided that the effect of height on the total hydraulic resistance 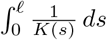 is negligible (which we argue to be the case in Appendix A2), this implies that a decrease of ψ_0_ by 1 Pa is equivalent to an increase in height by 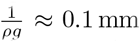 in terms of the impact on transpiration rate. The latter is closely linked to leaf photosynthesis and carbon assimilation (Damour et al. 2010). The hydraulic limitation hypothesis (Ryan and Yoder 1997) suggests that the closure of stomata due to leaf water potential decreasing with increasing tree height, and the associated decrease of carbon assimilation at the leaf level, induce the deceleration and eventual limitation of tree height growth. In cases where height growth is limited by hydraulic constraints, the above result may be used to quantify the extent of additional height growth that is enabled by an increase in soil water potential. Despite further supportive evidence (Hubbard et al. 1999, Martínez-Vilalta et al. 2007), Becker et al. (2000b) and Ryan et al. (2006) questioned the hydraulic limitation hypothesis. Among several alternative reasons suggested to play a more important role (Martínez-Vilalta et al. 2007), reduced turgor pressure has been hypothesised to limit height growth in terms of cell expansion and division (Fatichi et al. 2014). This variable is affected by low leaf water potential, induced by the same mechanism described in our model.

We made several simplifying assumptions in our model that can be relaxed while retaining the key mechanism. The steady-state assumption of the tree’s water balance is an idealisation since water storage in trees underlies diurnal fluctuations (Tyree and Zimmermann 2002). Nevertheless, in the long term, it may be a reasonable simplification. Linked to this is the non-consideration of the horizontal transport of water within the tree (Tyree and Zimmermann 2002) in our model. Although this is an important process, it plays a much smaller role than vertical movement, an observation that gave rise to the concept of hydraulic branch autonomy (Sprugel et al. 1991). We tacitly assumed the establishment of the *E*^*^-ψ(ℓ)^*^ equilibrium for given environmental conditions such as soil water potential ψ_0_ to occur instantaneously. Indeed, it has been shown that stomata open and close very quickly (Saliendra et al. 1995, Salleo et al. 2000, 2001). Nevertheless, applying the model to a much smaller time scale would require modifications e.g. in terms of a smooth change in stomata aperture over time in response to a changing environment, as well as a delay in its interaction with water potential. The balancing between the probability for embolism formation, which, in so-called vulnerability curves, is described as a function of water potential, and the repair of embolisms (Tyree and Zimmermann 2002) is only implicitly accounted for in our model, and could be considered in a mechanistic way. Lastly, while treated as a given exogenous parameter in our model, soil water potential is, in reality, subject to the functioning of the individual tree as well as the larger stand community.

## Acknowledgements

We are grateful for support of this project by a doctoral scholarship of the Heinrich-Böll Foundation. We also wish to thank Karl-Heinz Häberle for his helpful comments on this manuscript.

## Appendix

### A1 Explicit calculation of equilibrium leaf water potential ψ(ℓ)^*^

Let the simple algebraic sigmoid function

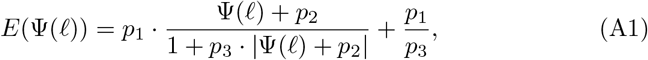

with parameters *p*_1_, *p*_2_, *p*_3_ > 0 controlling vertical dilatation, horizontal translation and steepness, respectively, describe the empirically measured response of transpiration rate E to leaf water potential ψ(ℓ). As opposed to certain alternative sigmoid function types such as the logistic function used in the regression by Tyree and Zimmermann (2002), this form allows the explicit calculation of the equilibrium leaf water potential in (5). It reads

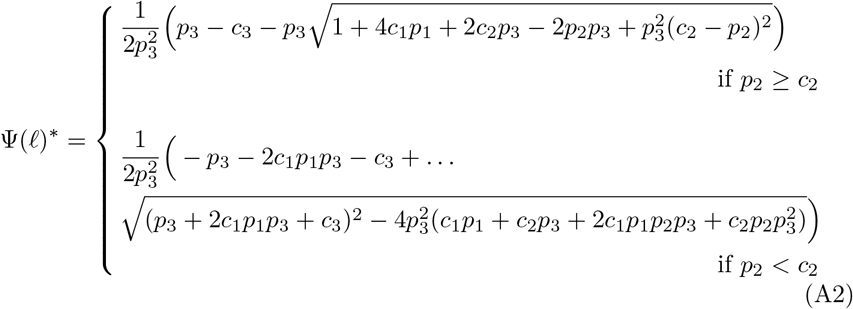

where *c*_1_ and *c*_2_ are as in (5), and where we abbreviated 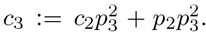 Inserting (A2) into (A1) yields the explicit form of the equilibrium transpiration rate, denoted f in (6).

### A2 Simplifying the total hydraulic resistance 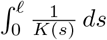

For shoot segments, the allometric relation *K*(*s*) = *a* ⋅ *C^b^*(*s*), where *C*(*s*) denotes the branch cross-sectional area at *s*, and *a* > 0, *b* > 1 are species-specific constants, is empirically documented for a range of species (Cochard et al. 2000, Cruiziat et al. 2002, Mencuccini 2002, Patino et al. 1995, Tyree et al. 1991, Yang and Tyree 1993, Zotz et al. 1998). This relationship illustrates the disproportionately strong impact of the smallest and final parts of the stembase-to-leaf pathway on the total hydraulic resistance, as opposed to a much smaller effect of thicker branches and the trunk. Empirical evidence for this was presented by Yang and Tyree (1993), Yang and Tyree (1994) and Tyree and Zimmermann (2002), who found that the major part of the water flow resistance in shoots was located in the leaf blade, petiole and smallest branches (in that order). Root hydraulic conductance has been studied less (Cruiziat et al. 2002, Tyree and Zimmermann 2002). Allometric relationships between cross-sectional area and conductance of root segment, similar to those for shoots, have been observed in some species (Doussan et al. 1999, Kotowska et al. 2015), although the allometric coefficients may differ from the ones for shoots (Gonzalez-Benecke et al. 2010). Analogous to the results for shoots, Tyree and Zimmermann (2002) found that the smallest parts of the roots, namely the radial non-vascular pathway from the surface of the fine roots to the vessels, as well as the fine roots themselves (in that order) account for most of the resistance in the root system.

These findings show that the shoot and root parts that dominate the total soil-to-leaf hydraulic resistance, namely the very smallest parts, are present in small and large trees alike, suggesting that soil-to-leaf hydraulic resistance depends little on the leaf’s position in the tree, the tree’s size or its age. This is in agreement with the theoretical considerations by West et al. (1999), Enquist (2000) and Becker et al. (2000a), who demonstrated that an appropriate tapering of the vascular conduits results in a hydraulic resistance that is independent of the path length. It may motivate to approximate the a priori tree-anatomy-specific soil-to-leaf resistance 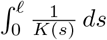 by a species-specific constant.

